# Extreme hot weather has stronger impacts on avian reproduction in forests than in cities

**DOI:** 10.1101/2020.01.29.924332

**Authors:** Ivett Pipoly, Bálint Preiszner, Krisztina Sándor, Csenge Sinkovics, Gábor Seress, Ernő Vincze, Veronika Bókony, András Liker

## Abstract

Climate change and urbanization are among the most salient human-induced changes affecting Earth’s biota. Extreme weather events can have high biological impacts and are becoming more frequent recently. In cities, the urban heat island can amplify the intensity and frequency of hot weather events. However, the joint effects of heat events and urban microclimate on wildlife are unclear, as urban populations may either suffer more from increased heat stress or become adapted to warmer temperatures. Here we test whether the effects of hot weather on reproductive success of great tits (*Parus major*) are exacerbated or dampened in urban environments compared to forest habitats. By studying two urban and two forest populations over six years, we show that 14-16 days-old nestlings have smaller body mass and tarsus length, and suffer increased mortality when they experience a higher number of hot days during the nestling period. The negative effects of hot weather on body mass and survival are significantly stronger in forests than in urban areas, where these effects are dampened or even reversed. These results suggest that urban birds are less vulnerable to extreme hot weather conditions than their non-urban conspecifics, possibly by adaptively evolving or flexibly adjusting nestling physiology to tolerate heat stress, and/or by adjusting parental behavior in response to heat. This finding highlights that endothermic vertebrates may be able to adapt to heat stress, which may help their populations cope with the joint challenges of climate change and urbanization.

**Significance statement:** Extreme weather events are becoming more frequent due to climate change and can have substantial effects on reproduction and survival of wild animals. Urban heat island can amplify the frequency of extreme hot weather events, making it potentially more harmful for city-dwelling organisms. Alternatively, urban populations living in warmer environments may adapt to better tolerate heat. We investigated these alternatives by comparing nestling development and survival between urban and forest great tit (*Parus major*) populations. We found that urban populations are less vulnerable to heat: nestling body mass and survival decreased rapidly with the increasing number of hot days in forests, while these effects were dampened in urban broods. Thus, urban populations may become adapted to better tolerate heat events.

## Introduction

Current large-scale environmental changes are substantially influencing the ecological conditions for Earth’s biota. Climate change is one of the dominant global processes that affect a wide range of organisms. Its effects include systematic changes in the long-term average meteorological conditions and also increases in the frequency of extreme weather events like heatwaves, droughts, and heavy storms [1–3]. These climatic changes are occurring in a world that is experiencing global-scale transformation of natural habitats into heavily modified anthropogenic habitats. For example, current estimates show that 77 % of the terrestrial habitats have already been modified by the direct effects of human activities such as agriculture and urbanization [4], and the remaining pristine habitats are also disappearing at a high rate [4–6] forcing an increasing number of wildlife populations to persist in anthropogenic environments [7]. Although the ecological and evolutionary consequences of different global processes such as climate change and anthropogenic land-use conversion are most often investigated separately, they obviously do not act independently. Thus to be able to better understand their current effects and to predict the changes likely induced in the future, we need to investigate their joint impacts in natural systems.

Hot days and heatwaves are among those extreme meteorological events that have become more frequent in the last few decades [2]. Extreme hot conditions can have strong biological impacts, affecting both survival and reproduction of organisms [8–10]. For example, the infamous heatwave that occurred in southern and western Europe in 2003 resulted in an estimated 70,000 heat-related human deaths [11] and was followed by detectable decreases in wild bird populations [12]. Similarly, extreme hot weather affects reproductive success negatively in a wide range of taxa, including both ectothermic [13, 14] and endothermic animals [10, 15–21]. These effects of extreme hot weather may be mediated by several, non-exclusive mechanisms, including both direct effects of heat stress on physiological and cognitive functioning [18, 20, 22] and indirect effects of heat *via* altered ecological conditions like food or water availability [23, 24].

Anthropogenic habitat change, especially urbanization, is likely to interact with extreme weather events because it fundamentally alters several basic environmental conditions, including microclimate and food availability [7, 25]. Urban heat island (UHI) refers to the generally higher ambient temperature in cities compared to the surrounding non-urban areas, which is largely generated by heat storage in buildings and sealed roads. Its intensity can be as high as + 5 °C (+ 9 °F) in some metropolitan areas [26–28], so organisms in cities experience higher temperatures and more frequent heat events compared to organisms in non-urban areas [28]. The UHI effect may interact with heat events in at least two ways. First, UHI can exacerbate the biological effects of extreme heat events. For example, models show that daytime temperature in temperate-zone cities during a heatwave can be higher by, on average, 2.8 °C as a result of the synergistic effect between UHI and heat events [29]. Moreover, heat stress in cities might be even higher during a heat event than the sum of the background UHI effect and the heatwave effect [30]. According to these model predictions, the number of hot days is higher in urban than in non-urban areas with an additional increase in UHI resulting in increased human mortality [30, 31]. Similarly, animals are more likely to reach their upper limit of thermal tolerance in cities during heat events than in non-urban areas [32]. Thus, urban populations may suffer a stronger reduction in survival and reproductive success than non-urban populations due to hot days (‘superimposed heat and UHI effects’ hypothesis).

Second, it is also possible that during long-term exposure to UHI, urban populations become adapted to higher temperatures (‘adaptation to UHI’ hypothesis). For example, heat tolerance, expressed as the thermal maximum where animals become unable to coordinate their motor performance, is hihger in urban compared to rural populations of acorn ants (*Temnothorax curvispinosus)* [33], water fleas (*Daphnia magna)*[34], and crested anoles *(Anolis cristatellus)* [35]. Moreover, survival and production of winged individuals in acorn ant colonies in an urban-rural reciprocal transplant experiment were higher in the home environment for both urban and rural colonies, probably due to local thermal adaptations [33, 36]. The reproductive advantage of heat-tolerant or heat-adapted phenotypes in hot urban conditions is also documented in the scale insect *Parthenolecanium quercifex*, as individuals collected from warm urban environments reproduce at a higher rate under hot than under cool laboratory conditions [37]. Although there is much less data on urban thermal adaptations in endotherms, the smaller body size of the rodent *Peromyscus maniculatus* [38] and reduced number of feathers in some birds [39] in urban populations might also reflect adaptations to the UHI.

So far, only a few studies have tested the above hypotheses for joint effects of extreme hot weather events and habitat urbanization. Their results show either additional mortality in cities in humans [30] or adaptation to UHI effects in water fleas [34]. Importantly, there is no field study for endotherm animal populations, despite that these are the most frequent targets of conservation efforts and play crucial roles in urban ecosystems. To start to fill this knowledge gap, here we tested whether heat events influence avian breeding success differently in cities and forest habitats. We analyzed six years of breeding biology data of two urban and two forest populations of great tits (*Parus major*), a small songbird that frequently breeds in both urban and forest habitats. In this species, similarly to several other birds, reproductive performance is typically lower in urban sites than in more natural areas [40, 41]. In the current study, we measured air temperature hourly at each site with own weather stations to calculate the number of hot days for each brood’s pre-fledging nestling period, and tested whether nestlings’ body mass, tarsus length, and survival are related to the number of hot days differently in urban and forest sites. According to the ‘superimposed heat and UHI effects’ hypothesis, we expect that heat events would result in stronger negative impacts on reproduction in urban than forest habitats. Alternatively, the ‘adaptation to UHI’ hypothesis predicts that heat events would have reduced impact in urban than forest populations.

## Results

### Habitat differences in heat events

Great tit offspring experienced markedly different temperature characteristics in urban and forest sites during the nestling season (Figure S1). Both the number of hot days (Figure S1A; urban-forest difference on log-odds scale: mean ± SE: 0.99 ± 0.13, t=7.39, df=750, p < 0.001) and the average temperatures within the nestling periods were significantly higher in the urban habitat (Figure S1B; urban-forest difference: 1.52 ± 0.27 °C, t=5.74, df=751, p < 0.001; model estimates controlled for year effects). That means the odds of hot days occurring during the nestling period was 2.7 times greater in the cities than in the forests: at least one hot day occurred in 179 out of 390 (45.9%) urban broods and in 77 out of 370 (20.8 %) forest broods. The highest maximum temperature was 40.1 °C in urban nestling periods (in June 2013) and 33.6 °C in forest nestling periods (in May 2014).

### Effects of hot days on reproductive success

#### Nestling body mass

In the simple model where predictors were the number of hot days, study site and their two-way interaction, the effect of hot days on nestling mass differed between sites (as indicated by a significant interaction between sites and number of hot days, Table 1) and between urban and forest habitats (according to the linear contrast post-hoc test comparing the effects between the two urban and the two forest populations, Table 2): increase in the number of hot days negatively affected nestling mass in forest broods while no such trend was present for the urban broods (Figure 1A). This result is consistent with the site-specific trends: the two forest populations had the most negative slopes for the relationship of nestling mass with the number of hot days, whereas the slope was flat or even positive in the two urban populations (Table S4).

**Table 1.** Relationship between the number of hot days during nestling development and nestlings’ body mass, tarsus length, and mortality, according to the simple models. Type 3 ANOVA (analyzis of deviance) table is shown for each model. Significant effects (p < 0.05) are highlighted in bold.

| Model terms | Nestling mass |  |  | Nestling tarsus length |  |  | Nestling mortality |  |  |
| --- | --- | --- | --- | --- | --- | --- | --- | --- | --- |
| | $\chi^2$ | df | P | $\chi^2$ | df | P | $\chi^2$ | df | P |
| Nr. hot days | 3.119 | 1 | 0.077 | **6.134** | **1** | **0.013** | **5.799** | **1** | **0.016** |
| Study site | **243.518** | **3** | **< 0.001** | **103.380** | **3** | **< 0.001** | **91.078** | **3** | **< 0.001** |
| Nr. hot days × study site | **29.289** | **3** | **< 0.001** | **9.555** | **3** | **0.023** | **11.844** | **3** | **0.008** |

For average nestling mass and tarsus length, the number of pairs was 535 and the number of broods was 674. For nestling mortality, the number of pairs was 600 and the number of broods was 760.

**Table 2.** Differences between urban and forest habitats in the effect of hot days on nestling’s body mass, tarsus length, and mortality. The table shows linear contrasts comparing the average slope of two urban *versus* two forest sites, calculated from the estimates of the simple models (left) and the most supported multi-predictor models (right) for each response variable. For nestling mass, results of two supported multi-predictor models are separated by “/”. Positive contrasts mean more positive slopes (i.e. less negative effects of hot days) in the urban habitat. Significant habitat differences (p < 0.05) are highlighted in bold.

| Response | Simple model |  |  |  | Multi-predictor model |  |  |  |
| --- | --- | --- | --- | --- | --- | --- | --- | --- |
| | contrast ± SE | df | t | p | contrast ± SE | df | t | p |
| Nestling mass | **0.338 ± 0.085** | **135** | **3.978** | **< 0.001** | **0.656 ± 0.166 / 0.720 ± 0.165** | **127 / 129** | **3.941 / 4.355** | **< 0.001 / < 0.001** |
| Nestling tarsus length | 0.046 ± 0.039 | 135 | 1.187 | 0.237 | 0.044 ± 0.075 | 127 | 0.590 | 0.556 |
| Nestling mortality* | **-0.408 ± 0.159** | **156** | **-2.564** | **0.010** | **-0.643 ± 0.281** | **149** | **-2.288** | **0.022** |

\* Estimates for nestling mortality are on the logit scale

**Figure 1.**
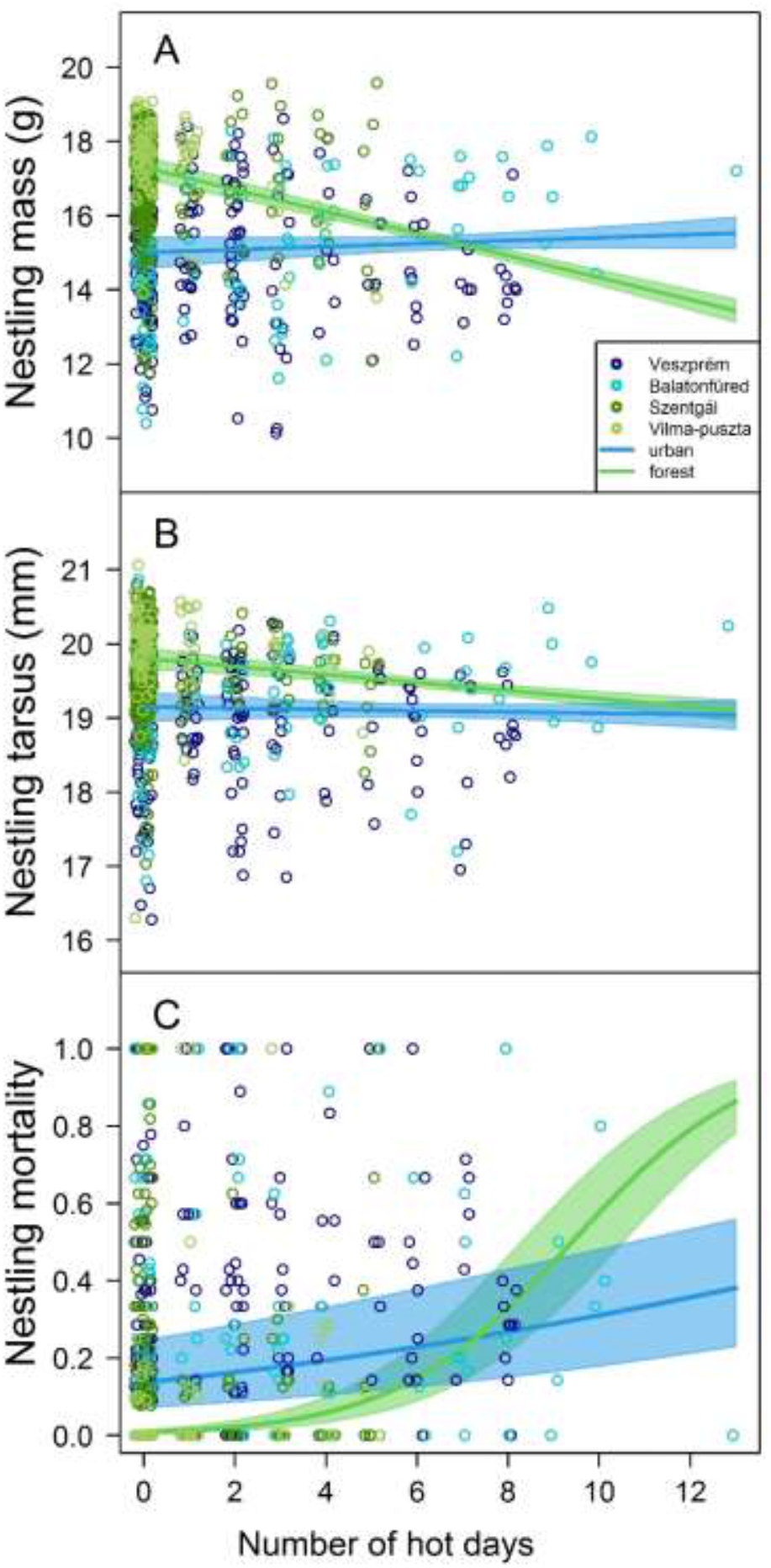
Relationships between the number of hot days during the nestling period and (A) body mass, (B) tarsus length, and (C) mortality of great tit nestlings. Circles represent brood means for nestling mass and tarsus length. Colored stripes show the 95% confidence band of the slope estimated from the simple models (Table S4).

We also tested whether the effect of hot days persist when further potentially important variables (i.e. hatching date, year, brood size an brood age) are taken into account. We built a set of models containing various combinations of these variables in addition to the number of hot days, study site and their interaction, and we compared these models using the information-theoretic approach based on AICc. Model selection for these multi-predictor models resulted in two supported models for nestling mass (ΔAICc < 2; the model with the third best fit had ΔAICc = 4; Table S1). The first supported model included year, brood age, and brood size besides the number of hot days, study sites, and their interaction; the second supported model contained the same predictors except brood size. Linear contrasts calculated from each of these supported multi-predictor models showed a significant overall habitat difference (Table 2) with a more negative effect of hot days in forest compared to urban populations (Table S4), corroborating the results of the simple model.

#### Nestling tarsus length

In the simple model, the interaction between site and number of hot days was significant (Table 1), due to the difference between the two urban sites, with a positive slope in one site and negative in the other (Table S4). The two forest sites both showed negative relationship between the number of hot days and nestling tarsus length (Table S4). The linear contrast for urban-forest difference suggested no systematic habitat difference in the effect of hot days on the tarsus length of nestlings (Table 2, Figure 1B).

Model selection for the multi-predictor model set resulted in a single supported model, which contained the number of hot days, study site, their interaction, the average temperature of the nestling period, brood age, and brood size (for the other models ΔAICc ≥ 6.68, Table S2). The result of linear contrast comparing urban and forest slopes from this model was qualitatively identical to the result from the simple model, showing no consistent habitat difference in the effect of hot days on nestling tarsus length (Table 2, Table S4).

#### Nestling mortality

The simple model showed that the effect of the number of hot days on nestling mortality differed significantly between study sites (Table 1). The slope of the relationship between nestling mortality and the number of hot days was positive in all populations, being highest in one of the forest sites and lowest in one of the urban sites (Table S4). The slopes also differed between urban and forest populations with a stronger effect in the latter (Table 2, Figure 1C), i.e. nestling mortality increased significantly more with the number of hot days in the forests than in the cities.

Multi-predictor model selection resulted in a single supported model, which was the full model of our model set (for the other models ΔAICc ≥ 42.87, Table S3), containing the number of hot days, study site and their interaction, the average temperature of the nestling period, year, hatching date, and the quadratic term of hatching date. The linear contrast calculated from this model showed a significantly steeper slope in forests compared to urban sites (Table 2), corroborating again the result of the simple model.

## Discussion

Our study found clear differences between urban and non-urban populations in the effects of hot days on important fitness components. Although the influence of thermal environment on fitness has been shown to vary with habitat urbanization in ectotherms [36], our study is the first to demonstrate such habitat-dependent responses to heat stress in an endothermic animal. While UHI caused a 1.19 °C higher temperature on average and there were more hot days in the cities compared to forests (Figure S1), hot days affected nestling mass and survival more negatively in the forest habitat type. These results are robust, as they are based on >600 broods from 6 years and four sites, and were qualitatively unchanged by taking into account several potentially confounding factors that may differ between urban and forest habitats, such as the start of breeding and brood size. These findings highlight that both lethal and sublethal effects of heat events (i.e. the proportion of surviving offspring and their quality in terms of body mass) can differ between urban and non-urban habitats. Although information about sublethal fitness costs of hot weather is scarce [9], developing under hot weather conditions may have long-term consequences for fitness. Our results thus support the ‘adaptation to UHI’ hypothesis and suggest that non-urban populations of birds (and possibly other vertebrates) are more vulnerable to extreme heat than urban populations.

Heat may affect the reproductive output of birds in at least two ways: directly through offspring physiology and indirectly through food availability. Firstly, the negative effects of hot days on offspring mass and survival may emerge due to the direct physiological consequences of heat stress. Nestlings usually cannot maintain stable body temperature in the first few days of their lives, and their metabolic processes also differ from those of adults [42] leaving them potentially more vulnerable to extreme heat than adults. The heat responses of dependent offspring are poorly known, although in birds the nestling phase seems to be the most relevant period during ontogeny regarding heat effects compared to the incubation and post-fledging periods [43]. In hot environments, individuals can less effectively dissipate the excess heat as a consequence of increased metabolic rate, leading to hyperthermia. Increasing metabolism causes decreased utilization of food and faster mobilization of energy reserves, thus it may cause lower body mass. Additionally, during heat stress, evaporative water loss is elevated as the organism tries to cool itself by evaporation to maintain body temperature in the physiologically normal range, causing dehydration [44, 45]. Thus, increased metabolic rate and water loss both can lead to decreased body mass, as it has been shown in several bird species [16, 46–48].

These direct physiological effects of heat might be better tolerated by urban animals due to changes in their physiological and morphological characteristics. For example, we found in the same study populations that urban great tit nestlings have fewer feathers and increased bare body surfaces than forest nestlings [39], which may facilitate heat dissipation. Reduced body size may also help to cope with the heat in cities because animals with smaller size have a higher surface-biomass ratio which facilitates heat loss. Bergmann’s rule [49] predicts that reduced size is beneficial against dehydration and overheating in warmer environments [44] (but see [50]) such as cities. This idea is supported by urban-rural comparisons of invertebrate communities [51], as well as by a study of water flea populations which found that urban individuals were generally smaller than rural ones, and smaller individuals were more heat tolerant [34]. In birds, a study of white-browed scrubwrens (*Sericornis frontalis*) also found that smaller individuals survived better when more extreme hot and dry events occurred [52]. Urban birds are usually smaller than their non-urban conspecifics [53–55] and this is also the case with the great tits in our study populations [40]. Although experimental evidence shows that low availability of nestling food is a major cause of the reduced size of urban great tit nestlings [56], selection for better heat tolerance might also contribute to it. Finally, life-history changes may also help animals to cope with the direct effects of urban heat. For example, urban birds usually have lower clutch size than forest birds [40, 53] which may also be advantageous for heat dissipation when environmental temperature is high because fewer offspring produce less heat and have more space to maintain distance in the nest, which helps better heat conduction.

Secondly, indirect effects may also contribute to lower body mass and higher mortality in response to heat events. For example, hot weather may reduce food availability for insectivorous nestlings by affecting their parents and/or their prey. Caterpillars are the main source of nestling diet in great tits and many other birds, and increasing temperature strongly decreases the time to pupation in several lepidopteran species [57–59], so optimal caterpillar food may be available for a shorter time in hot periods. Moreover, the growth of caterpillars declines rapidly above a critical temperature [58] and their mortality increases when temperature is constantly high [59, 60]. These heat effects may not only result in less food for nestlings but may also contribute to dehydration because caterpillars are the most water-rich items in the diet of nestlings [61]. These effects might be stronger in forests where the amount of available caterpillar prey is much higher in general [40] and the vast majority of nestling diet consists of caterpillars. In such environments, it might be more difficult for parent birds to compensate for a reduction in caterpillar biomass compared to cities, where nestlings are usually fed a greater proportion of other food types [62, 63]. Additionally, the foraging activity of parents may also decrease when temperature is high [16, 64, 65], resulting in less food delivered to the offspring, and this effect might also be diminished in urban populations if parent birds have adapted to tolerate heat better (see above). Further studies are required to test whether heat affects parental care differentially in urban and non-urban habitats. Our study suggests that the increasing frequency of hot days expected as a consequence of ongoing climate change would be more harmful for natural than urban populations, further endangering biodiversity. For example, our results predict that the reproductive advantage of non-urban populations (e.g. higher body mass and lower mortality of nestlings) over urban populations can be reversed when the number of hot days experienced by nestlings becomes high (exceeds 7-8 days in our study). This raises the intriguing possibility that urban populations might serve as sources for the conservation of some species in the future. Adaptation to constantly warmer urban environments may be achieved by microevolution, epigenetic changes, and/or phenotypic plasticity [66]. In some ectotherm species, artificial thermal selection and common garden experiments have revealed that one or more of these mechanisms can play a role in the increased upper thermal tolerance of urban populations [34–36, 67]. To our knowledge, no such experiments have yet been done to identify the mechanisms of higher thermal tolerance in urbanized birds or other endothermic animals, which is an important knowledge gap in our understanding of urban adaptations. Therefore, there is a great need for further studies that explore the differences in thermal tolerance between different habitats to understand how wild animals are affected by, and may adapt to, the interacting challenges of climate change and urbanization in the Anthropocene.

## Methods

### Data collection

We installed nest-boxes for great tits in an urban site (city of Veszprém 47°05’17”N, 17°54’29”E), and in two sites in natural habitats (forests near Vilma-puszta 47°05’06.7”N, 17°51’51.4”E and Szentgál 47°06’39”N, 17°41’17”E) in 2012, and additionally in another urban site (Balatonfüred 46°57’30”N, 17°53’34”E) in 2013. Urban nest-boxes are located mostly in public parks, university campuses, and a cemetery, where vegetation contains both native and introduced plant species. Forest study sites are located in deciduous woodlands, characterized by beech *Fagus sylvatica* and hornbeam *Carpinus betulus* (in Szengtál) or downy oak *Quercus cerris* and South European flowering ash *Fraxinus ornus* (in Vilma-puszta). See detailed description of study sites in [40]. In March 2013, we installed a WH 2080 weather station (Ambient, LLC, AZ, USA) at each study site that recorded hourly temperature (°C) data throughout the six years of the study.

Great tits usually raise one or two broods per breeding season. We collected data on all breeding attempts at each site from 2013 to 2018, by recording the number of eggs and nestlings in the nest-boxes every 3-4 days from March to the end of July. We captured parent birds using a nest-box trap 6-15 days after their first nestling had hatched. We determined parents’ sex based on their plumage characteristics and banded each bird with a unique combination of a numbered metal band and three plastic color bands. Breeding adult birds that had been color-banded on previous occasions were identified from recordings made during the nestling period by using a small, concealed video camera attached to the nest-boxes [68]. In these video samples, we considered a color-banded individual to be a parent bird if it was recorded to enter the nest-box with food at least once. Close to fledging (at day 14-16 post-hatch; day 1 being the hatching day of the first nestling in the brood), we measured nestlings’ body mass with a Pesola spring balance (± 0.1 g) and tarsus length with a Vernier caliper (± 0.1 mm).

All procedures applied during our study were in accordance with the guidelines for animal handling outlined by ASAB/ABS (www.asab.org) and Hungarian laws. We have all the required permissions for capturing, measuring of the birds and monitoring their breeding from the Government Office of Veszprém County, Nature Conservation Division (former Middle Transdanubian Inspectorate for Environmental Protection, Natural Protection and Water Management; permission number: 24861/2014 and VE-09Z/03454-8/2018) and from the National Scientific Ethical Committee on Animal Experimentation (permission number: VE/21/00480-9/2019).

### Data processing

We used both the first and second annual broods of each pair in the study. We omitted broods where we could not precisely identify the nestling period due to unknown dates of hatching and/or the death of the entire brood before hatching, and also those broods where we had gaps in the temperature data for more than two days during the nestling period (n= 190 broods). For analyzing the effects of weather on nestling size, we used the average body mass and the average tarsus length of each brood as response variables, and we included broods where at least one offspring was alive at the time of nestlings’ measuring and banding (i.e. the age of 14-16 days). Because chick size varies with age, we omitted those few broods where the nestlings were measured before or after 14-16 days of nestling age (n= 30); thus we had n= 674 broods for nestling size analyzes.

For analyzing the effects of weather on nestling mortality, we used broods in which at least one offspring was alive on the third day after the hatching of the first nestling. We excluded a small number of broods (n= 7) that failed within the first three days after hatching because the average interval between our nest monitoring visits was three days, so estimating the number of hatched chicks and their survival before the first nest check after hatching would be very uncertain. We also excluded those broods where chick mortality was likely unrelated to weather conditions (n= 7), i.e. occurred due to predation or human disturbance (i.e. recorded during the next observation after the occasion of capturing the parent bird on its nest). This yielded n= 760 broods for nestling mortality analyzes, from which 385 had no chick mortality, 319 had partial mortality, and 56 had complete brood loss during the nestling period.

We calculated all weather variables (average temperature and number of hot days, detailed below) for the nestling period of each brood individually. For investigating nestling size, weather variables were calculated for days from the day of hatching of the first chick to the day preceding the day of banding and measuring the nestlings. This sample included only broods where banding occurred at age 14-16 days (mean ± SE: 14.04 ± 0.03). For analyzing nestling mortality, we calculated weather variables for the period from the day of hatching of the first offspring to either the day preceding the day of recording the last nestling’s death or the day preceding the day of banding and measuring the nestlings (mean ± SE: 13.72 ± 0.07, range: 3 – 18 days). We calculated the average temperature as the mean of hourly temperatures over the nestling period. There are several methods for quantifying extreme hot weather [69], using various thresholds from 21 to 43 °C (70 to 110 °F) depending on the geographic location. We calculated the number of hot days during each nestling period using a threshold specific to our region. To calculate a threshold for hot days, first, we defined a reference period from the earliest hatching date to the latest chick banding date recorded in our six-year data set (pooling all sites and years), which ranged from 9th April to 15th July. Then, to estimate the typical temperatures in our geographic area during this reference period, we used a 26-years dataset from an external reference weather station located in Szentkirályszabadja, a small village near Veszprém (47°57’06” É, 17°58’10” K, ca. 9.5 – 22 km from our study sites). This weather station is maintained by the Hungarian Meteorological Service and its temperature data are available from the National Oceanic and Atmospheric Administration database (www.ncdc.noaa.gov), with records of air temperature every three hours a day (0, 3, 6, 12, 15, 18, 21 UTC) since 1993. Using this temperature dataset, we calculated the 90 % percentile of all daily maximum temperatures for the reference period from this 26-year long dataset, which was 28.7 °C. Finally, we calculated the number of hot days as the number of days when the daily maximum temperature was higher than the 28.7 °C threshold during the nestling period for each brood. Fixed temperature thresholds for calculating hot days are widely used in ecological and evolutionary research [69], based on the assumption of a fixed threshold for heat tolerance in adults and offspring [18, 19, 52, 70].

### Statistical analyzes

Analyzing the effects of extreme weather is challenging. Extreme events are rare by definition, so their distribution is strongly skewed. Furthermore, weather variables are often correlated with each other and with other variables influencing reproductive success (e.g. with calendar date), so multicollinearity can be a problem in models containing multiple predictors. Statistical methods that can handle multicollinearity, such as covariance-based structural equation modelling, are less well suited for handling non-normal residual distributions and the non-independence structure of ecological data (e.g. multiple broods per pair). Therefore, here we used general and generalized linear mixed-effects (LME and GLMM) models and Akaike’s information-criterion (AIC) based model selection [71] to focus on the interaction between habitat type and the number of hot days in explaining reproductive variables. Mixed-effects models can be applied appropriately to non-normal and non-independent data, while model selection based on the models’ total explanatory power (model fit) is not sensitive to multicollinearity [72].

All analyzes were run in R (version 4.0.0) [73]. First, to examine the temperature characteristics of our study sites, we compared the number of hot days and the average temperature in the nestling periods (calculated separately for each brood) between urban and forest habitats. We constructed a generalized linear model with generalized Poisson distribution for the number of hot days using “glmmTMB” package [74] to handle overdispersion, and a linear model for the average temperatures. In both models, study site (4 sites) and year (6 years) were the predictors. To statistically compare the mean of each meteorological variable between the two habitat types, we calculated marginal means from the models for each study site, then calculated a linear contrast for habitat comparison as the difference between the average of the two urban sites versus the average of the two forest sites, using the “emmeans” R package [75]. We used this approach rather than including habitat type as a fixed effect and site as a random effect in the models because variance estimations of random effects with few levels are unreliable [76, 77], whereas including both habitat type and site as fixed effects would have resulted in strong collinearity between these two factors [78]. Instead, we treated the four sites as if they were two control groups and two treatment groups in an experiment, and we used a pre-planned comparison to test the prediction that the two treatment (i.e. urban) groups would differ from the two control (i.e. forest) groups. Such pre-planned comparisons are a powerful approach for testing *a priori* hypotheses [79].

To analyze average nestling mass and average tarsus length as response variables, we used the “lme” function of the “nlme” R package [80] which assumes Gaussian error. For nestling mortality (proportion of hatched chicks that died by the age of 14-16 days) as the response variable, we used GLMM models using the “glmer” function of the “lme4” R package [81] with binomial error distribution and logit link function. All of our models contained pair identity as a random factor to control for the non-independence of broods produced by the same pair. Broods got the same pair ID if both the female and the male parents were the same, and a new pair ID was assigned to the brood when it had at least one different parent. There were 111 pairs of parents that had more than one brood in our dataset, ranging 2-6 broods/pair. Additionally, to handle overdispersion in models of nestling mortality, we used an observation-level random effect [82] and ran the models with the BOBYQA optimizer [81].

First, we built a simple model for each of the three response variables (nestling mass, tarsus length, and mortality) including only the number of hot days (used as a numeric covariate) and study site (used as a factor with four levels), and their two-way interaction as predictors. Then, to statistically compare the effect of hot days on nestling size and mortality between the two habitat types, we calculated the slopes of the relationship between reproductive success measurements and the number of hot days for each study population, and we compared the average of the two urban slopes with the average of the two forest slopes using a linear contrast.

To infer the robustness of the results from the simple models, we also built multi-predictor models which included further potentially important predictor variables. In all models of our three response variables, we included year as a categorical variable, the average temperature during the nestling period and hatching date as numeric covariates, the latter defined as the number of days elapsed from the 1^st^ January annually to the hatching of the first nestling in each brood. We also added the quadratic term of hatching date, because a preliminary inspection of diagnostic graphs suggested the possibility of non-linear seasonal changes in the response variables. For models of nestling size (i.e. average body mass and average tarsus length of nestlings), we additionally included brood age (i.e. the number of days elapsed from the hatching of the first chick to the day of nestlings’ measuring, ranging from 14 to 16 days), and brood size (i.e. the number of offspring alive at measuring). We standardized all numeric predictors using z-transformation for our multi-predictor analyzes to avoid model convergence problems due to different scales of variables. Variance inflation factor (VIF), a measure of multi-collinearity, ranged from 1.04 to 6.24 for the predictors in these multi-variate models. Because multi-collinearity with VIF values > ca. 2 can lead to unreliable standard errors, non-significant parameter estimates from such models must be treated with caution [72]. However, the set of predictors that yields the best explanatory power can be correctly identified by model selection based on model fit statistics (like AIC), even if multi-collinearity is present [71]. Therefore, we created a candidate model set for each response variable using all combinations of predictors but always including the number of hot days, study site, and their interaction (because this interaction was the focus of our study, and our goal with the model-selection procedure was to see if this interaction yields the same habitat contrasts in the multi-predictor models as alone in the simple models). The model set contained 33 models for nestling size (Table S1-S2) and 9 models for nestling mortality (Table S3). We compared the models in each model set based on their AICc (AIC corrected for sample size) to identify the most supported model(s) for each response variable [71] using the “model.avg” function of the “MuMIn” R package [83]. From the model(s) with superior support based on ΔAICc and Akaike weight, we calculated the slope for each study site and the linear contrast that compares the effect of hot days between urban and forest sites the same way as from the simple models.

## Acknowledgements

We thank several field assistants for helping during fieldwork. The project was primarily financed by the National Research, Development and Innovation Office of Hungary (NKFIH, grants K84132, K112838 & K132490 to AL), and by the Hungarian Academy of Sciences. It was also supported by the TKP2020-IKA-07 project financed under the 2020-4.1.1-TKP2020 Thematic Excellence Programme by NKFIH, and by the European Union with the co-funding of the European Social Fund (TÁMOP-4.2.2.A-11/1/KONV-2012-0064). I.P. was supported by the ÚNKP-20-4 New National Excellence Programme of the Ministry of Innovation and Technology from the Source of the National Research, Development and Innovation Fund. V.B. was supported by the János Bolyai Scholarship of the Hungarian Academy of Sciences. EV was supported by the PD-134985 grant by NKFIH and by the MSCA EF Seal of Excellence IF-2019 grant by Vinnova, the Swedish Governmental Agency for Innovation Systems (grant number: 2021-01102). K.S. was supported by MTA-ELTE Comparative Ethology Research Group (F01/031).

## Supplementary Material for

**Figure S1.**
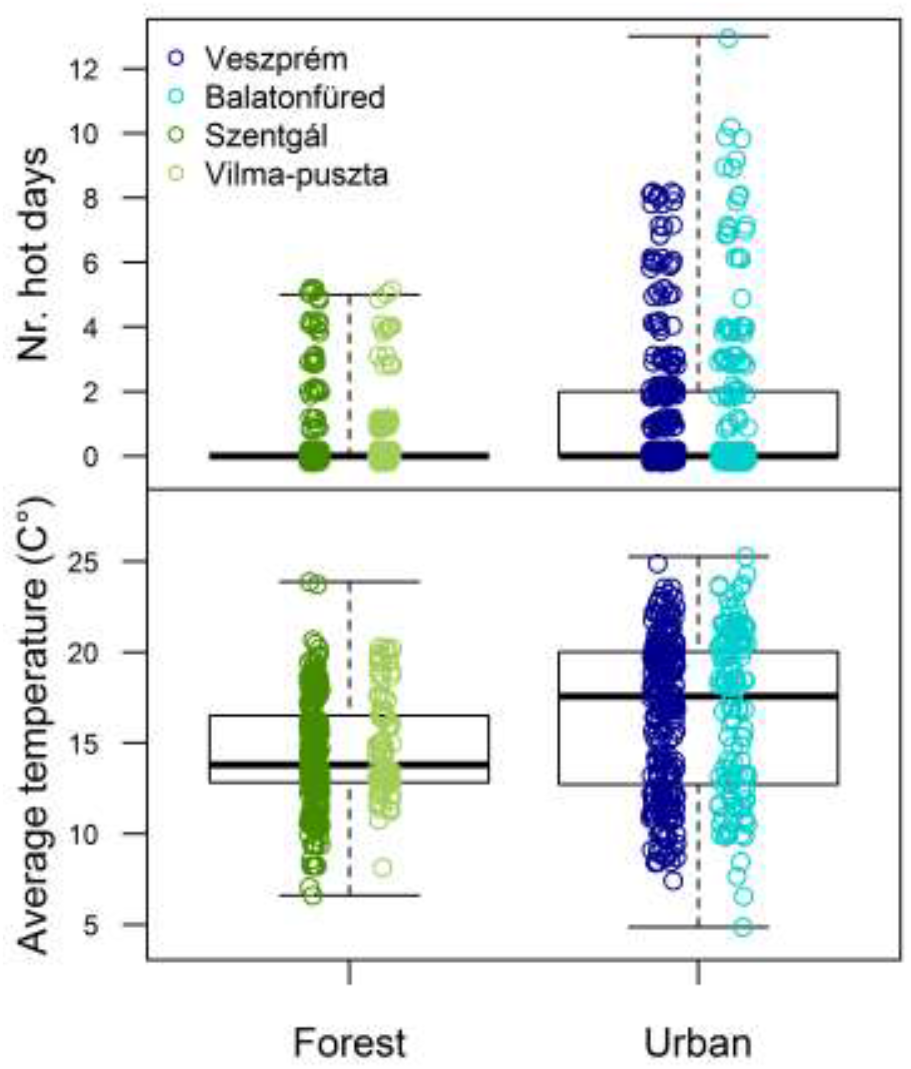
Distribution of the number of hot days and average temperature during the nestling periods in our study sites. Boxes show the interquartile range, the thick line is the median, and whiskers refer to the range of data distribution. Data points represent great tit broods (green: forest, blue: urban).

**Table S1.** Result of AICc-based model selection for average nestling mass. Supported models are highlighted in bold.

| Model | df | logLik | AICc | ΔAICc | Akaike weight |
| --- | --- | --- | --- | --- | --- |
| **456789** | **17** | **-1220.19** | **2475.31** | **0** | **0.55** |
| **45789** | **16** | **-1221.97** | **2476.78** | **1.47** | **0.26** |
| 23456789 | 19 | -1220.17 | 2479.49 | 4.19 | 0.07 |
| 1456789 | 18 | -1221.48 | 2480.00 | 4.69 | 0.05 |
| 145789 | 17 | -1223.37 | 2481.67 | 6.36 | 0.02 |
| 2345789 | 18 | -1222.4 | 2481.84 | 6.53 | 0.02 |
| 123456789 | 20 | -1221.12 | 2483.52 | 8.22 | 0.01 |
| 46789 | 16 | -1226.23 | 2485.29 | 9.99 | 0 |
| 12345789 | 19 | -1223.17 | 2485.51 | 10.2 | 0 |
| 4789 | 15 | -1227.84 | 2486.41 | 11.11 | 0 |
| 2346789 | 18 | -1226.43 | 2489.91 | 14.61 | 0 |
| 146789 | 17 | -1227.61 | 2490.15 | 14.84 | 0 |
| 234789 | 17 | -1227.96 | 2490.85 | 15.54 | 0 |
| 14789 | 16 | -1229.11 | 2491.05 | 15.75 | 0 |
| 2345679 | 14 | -1232.31 | 2493.25 | 17.95 | 0 |
| 12346789 | 19 | -1227.39 | 2493.94 | 18.63 | 0 |
| 45679 | 12 | -1234.77 | 2494.02 | 18.71 | 0 |
| 1234789 | 18 | -1228.84 | 2494.72 | 19.42 | 0 |
| 12345679 | 15 | -1232.37 | 2495.48 | 20.17 | 0 |
| 145679 | 13 | -1235.07 | 2496.68 | 21.38 | 0 |
| 4679 | 11 | -1241.41 | 2505.22 | 29.91 | 0 |
| 234679 | 13 | -1239.51 | 2505.57 | 30.26 | 0 |
| 1234679 | 14 | -1239.99 | 2508.62 | 33.31 | 0 |
| 14679 | 12 | -1242.58 | 2509.63 | 34.32 | 0 |
| 1234579 | 14 | -1245.06 | 2518.77 | 43.46 | 0 |
| 4579 | 11 | -1249.02 | 2520.44 | 45.13 | 0 |
| 14579 | 12 | -1249.51 | 2523.49 | 48.18 | 0 |
| 234579 | 13 | -1248.72 | 2524.00 | 48.69 | 0 |
| 123479 | 13 | -1251.77 | 2530.09 | 54.78 | 0 |
| 479 | 10 | -1255.36 | 2531.04 | 55.74 | 0 |
| 23479 | 12 | -1254.36 | 2533.20 | 57.89 | 0 |
| 1479 | 11 | -1256.61 | 2535.61 | 60.31 | 0 |
| (Null) | 3 | -1369.13 | 2744.30 | 269 | 0 |

Model term codes: 1=average temperature of nestling periods, 2=(hatching date)<sup>2</sup>, 3=hatching date, 4=number of hot days, 5=brood age, 6=brood size, 7= study site, 8=years, 9=number of hot days × study site. Null model contains only the random effect on nestlings' body mass.

**Table S2.**
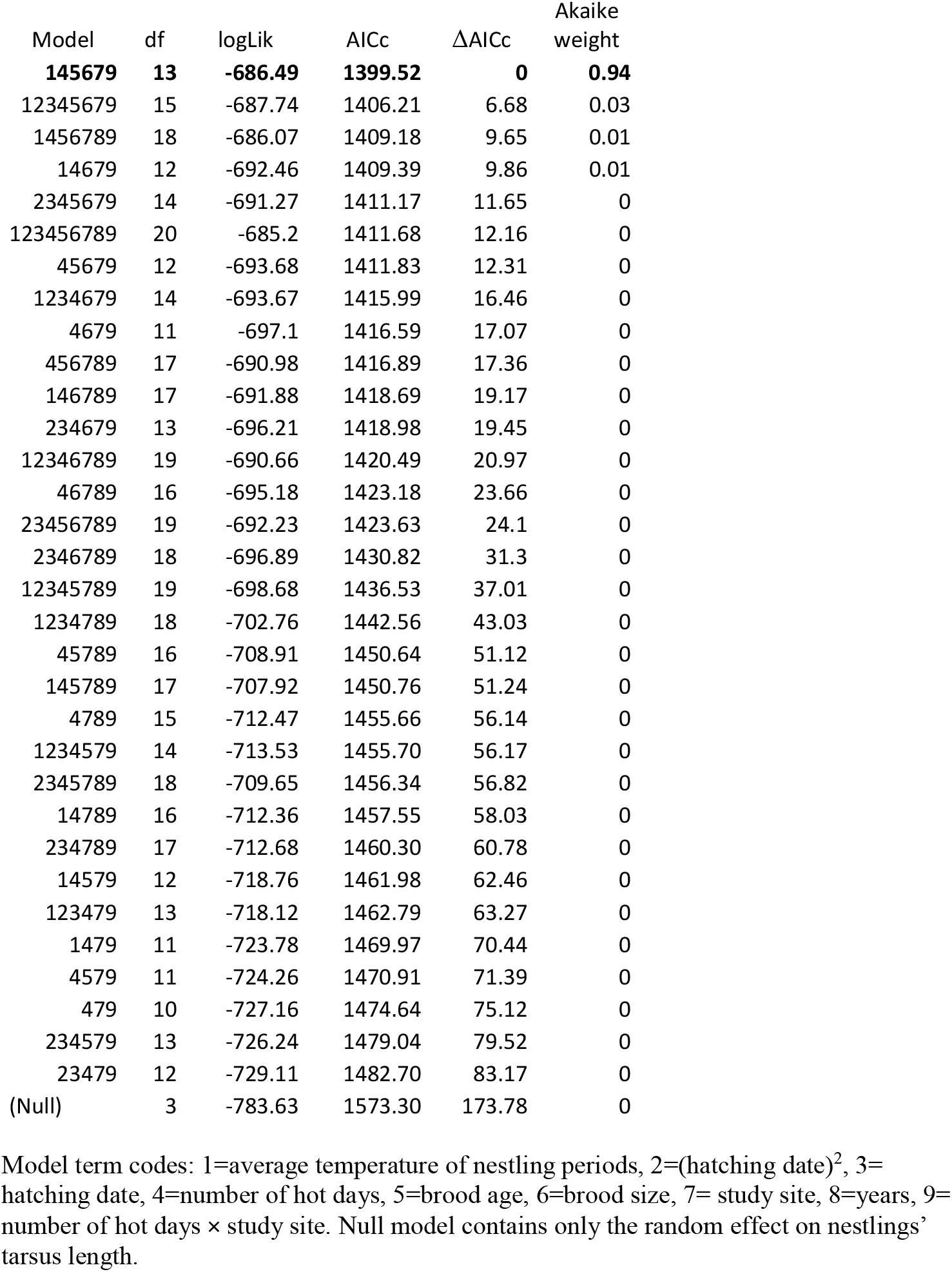
Result of AICc-based model selection for average nestling tarsus length. Supported model is highlighted in bold.

| Model | df | logLik | AICc | ΔAICc | Akaike weight |
| --- | --- | --- | --- | --- | --- |
| **145679** | **13** | **-686.49** | **1399.52** | **0** | **0.94** |
| 12345679 | 15 | -687.74 | 1406.21 | 6.68 | 0.03 |
| 1456789 | 18 | -686.07 | 1409.18 | 9.65 | 0.01 |
| 14679 | 12 | -692.46 | 1409.39 | 9.86 | 0.01 |
| 2345679 | 14 | -691.27 | 1411.17 | 11.65 | 0 |
| 123456789 | 20 | -685.2 | 1411.68 | 12.16 | 0 |
| 45679 | 12 | -693.68 | 1411.83 | 12.31 | 0 |
| 1234679 | 14 | -693.67 | 1415.99 | 16.46 | 0 |
| 4679 | 11 | -697.1 | 1416.59 | 17.07 | 0 |
| 456789 | 17 | -690.98 | 1416.89 | 17.36 | 0 |
| 146789 | 17 | -691.88 | 1418.69 | 19.17 | 0 |
| 234679 | 13 | -696.21 | 1418.98 | 19.45 | 0 |
| 12346789 | 19 | -690.66 | 1420.49 | 20.97 | 0 |
| 46789 | 16 | -695.18 | 1423.18 | 23.66 | 0 |
| 23456789 | 19 | -692.23 | 1423.63 | 24.1 | 0 |
| 2346789 | 18 | -696.89 | 1430.82 | 31.3 | 0 |
| 12345789 | 19 | -698.68 | 1436.53 | 37.01 | 0 |
| 1234789 | 18 | -702.76 | 1442.56 | 43.03 | 0 |
| 45789 | 16 | -708.91 | 1450.64 | 51.12 | 0 |
| 145789 | 17 | -707.92 | 1450.76 | 51.24 | 0 |
| 4789 | 15 | -712.47 | 1455.66 | 56.14 | 0 |
| 1234579 | 14 | -713.53 | 1455.70 | 56.17 | 0 |
| 2345789 | 18 | -709.65 | 1456.34 | 56.82 | 0 |
| 14789 | 16 | -712.36 | 1457.55 | 58.03 | 0 |
| 234789 | 17 | -712.68 | 1460.30 | 60.78 | 0 |
| 14579 | 12 | -718.76 | 1461.98 | 62.46 | 0 |
| 123479 | 13 | -718.12 | 1462.79 | 63.27 | 0 |
| 1479 | 11 | -723.78 | 1469.97 | 70.44 | 0 |
| 4579 | 11 | -724.26 | 1470.91 | 71.39 | 0 |
| 479 | 10 | -727.16 | 1474.64 | 75.12 | 0 |
| 234579 | 13 | -726.24 | 1479.04 | 79.52 | 0 |
| 23479 | 12 | -729.11 | 1482.70 | 83.17 | 0 |
| (Null) | 3 | -783.63 | 1573.30 | 173.78 | 0 |

Model term codes: 1=average temperature of nestling periods, 2=(hatching date)<sup>2</sup>, 3=hatching date, 4=number of hot days, 5=brood age, 6=brood size, 7= study site, 8=years, 9=number of hot days × study site. Null model contains only the random effect on nestlings' tarsus length.

**Table S3.** Result of AICc-based model selection for nestling mortality. Supported model is highlighted in bold.

| model | df | logLik | AICc | ΔAICc | Akaike weight |
| --- | --- | --- | --- | --- | --- |
| **1234567** | **18** | **-1032.7** | **2102.33** | **0** | **1** |
| 234567 | 17 | -1055.19 | 2145.20 | 42.87 | 0 |
| 123457 | 13 | -1074.94 | 2176.37 | 74.04 | 0 |
| 4567 | 15 | -1088.84 | 2208.32 | 105.99 | 0 |
| 14567 | 16 | -1088.53 | 2209.79 | 107.46 | 0 |
| 23457 | 12 | -1105.89 | 2236.20 | 133.87 | 0 |
| 1457 | 11 | -1125.98 | 2274.31 | 171.98 | 0 |
| 457 | 10 | -1130.95 | 2282.19 | 179.86 | 0 |
| (Null) | 3 | -1203.8 | 2413.62 | 311.29 | 0 |

Model term codes: 1=average temperature of nestling periods, 2=(hatching date)<sup>2</sup>, 3=hatching date, 4=number of hot days, 5= study site, 6=years, 7= number of hot days × study site. Null model contains only the random effect on nestling mortality.

**Table S4.**
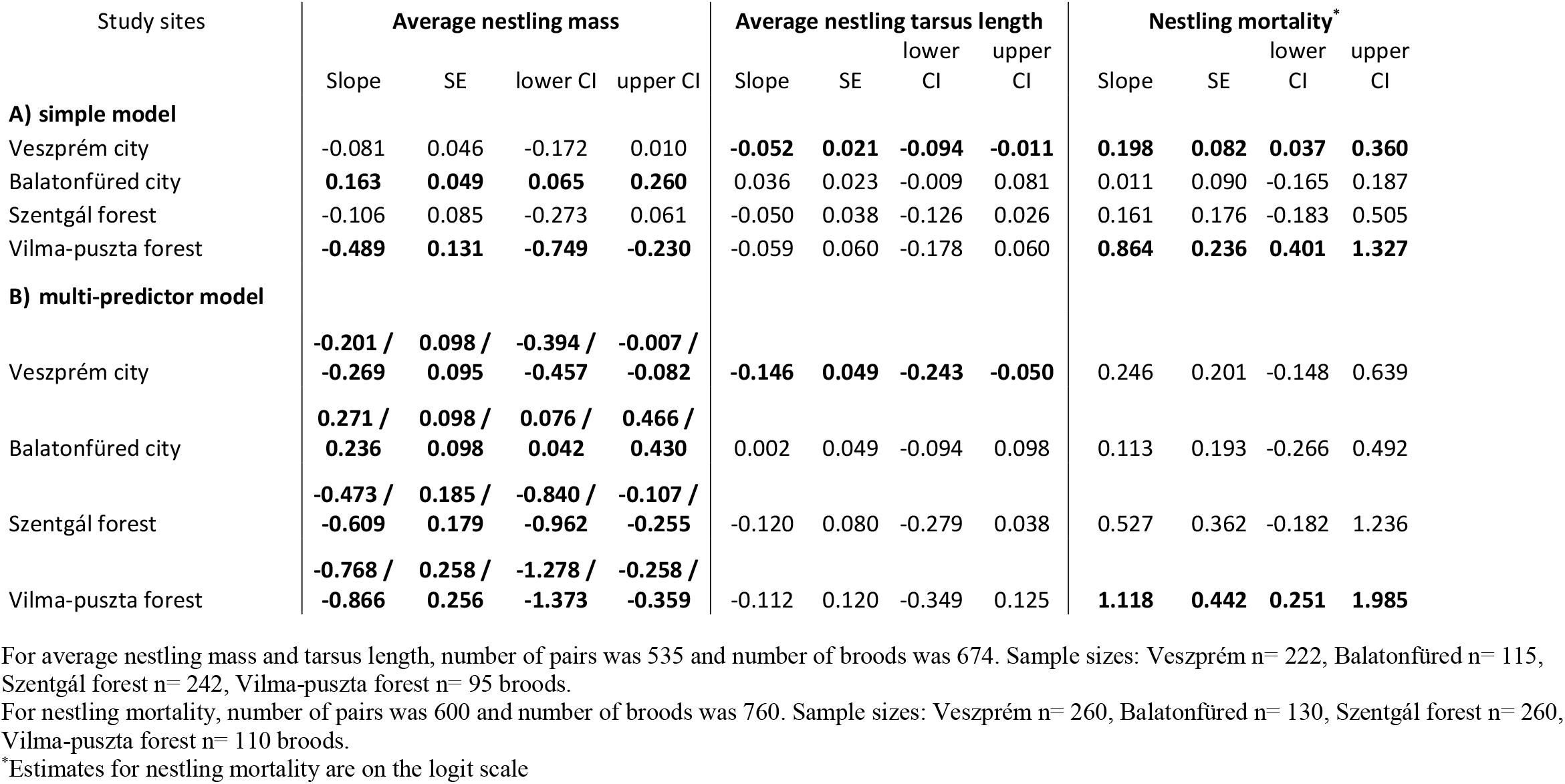
The effect of the number of hot days during the nestling period on reproductive success in great tits. The slope of the relationship with 95% confidence interval (CI) is shown for each site, as estimated by marginal means from A) the simple models (containing number of hot days, study site, and their two-way interaction) and B) the supported multi-predictor models (Table 1). For nestling mass, results of the two supported multi-predictor models are separated by “/”. Slopes significantly different from zero (i.e. zero not included between the lower and upper limit of CI) are highlighted in bold.

## Notes

**Competing interest statement:** The authors declare no competing interests

### Competing Interest Statement

The authors have declared no competing interest.

